# Corticotropin-releasing factor distribution in the brain of the brown anole lizard

**DOI:** 10.1101/2021.02.16.431399

**Authors:** David Kabelik

**Affiliations:** Department of Biology, Rhodes College, Memphis, TN, 38112 USA; Program in Neuroscience, Rhodes College, Memphis, TN, 38112 USA

**Keywords:** corticotropin-releasing factor, corticotropin-releasing hormone, neural distribution, lizard, reptile

## Abstract

Corticotropin-releasing factor (CRF) is best known for its involvement in peripheral glucocorticoid release across vertebrate species. However, CRF is also produced and released throughout various brain regions to regulate central aspects of the stress response. While these various CRF populations have been described extensively in mammals, less is known about their distributions in other amniotes, and only a handful of studies have ever examined CRF distributions in reptiles. Out study is the first to map CRF cell and fiber distributions in the brain of a lizard, the brown anole (*Anolis sagrei*). Our results indicate that brown anole CRF distributions are highly similar to those in snakes and turtles. However, unlike in these other reptile species, we find immunofluorescent CRF neurons in a few additional brown anole locations, most notably the supraoptic nucleus. The CRF distribution in the present study is also similar to published CRF descriptions in mammals and birds, although our findings, as well as the other published reports in reptiles, collectively suggest that reptiles possess a slightly more restricted distribution of CRF cell populations than do mammals and birds.

## Introduction

Corticotropin-releasing factor (CRF), also known as corticotropin-releasing hormone, is best known for causing the release of adrenocorticotropic hormone from the anterior pituitary gland, leading to eventual glucocorticoid release – a major component of an organism’s stress response and the hypothalamic-pituitary-adrenal axis [1]. In nonmammalian vertebrates, CRF also possesses an additional role, in that its release into the hypophyseal portal blood not only releases ACTH from the anterior pituitary, but it also acts as a thyrotropin-releasing factor, thereby exerting additional metabolic effects on the organism [2]. However, numerous central populations of CRF-releasing neurons also exist across vertebrates, and many project centrally rather than being released into the hypophysial portal blood [2]. These centrally projecting neurons seem to play a neuromodulatory role in central stress signaling and responsiveness, including the mediation of cognitive and emotional aspects of the stress response [1].

Distributions of CRF-producing neuron populations have been described in various mammalian [3–6] and avian [7] taxa. However, few descriptions of central CRF distributions have been published in reptiles. Reptiles, a highly understudied vertebrate lineage [8], are an important amniote group that provides opportunities for comparison with mammals and birds. Previously reported examinations of CRF distribution in reptiles have been exclusively in snakes and turtles [9–13], with no published distributions any lizard species.

In the present study, we use immunohistochemistry to examine the distribution of CRF-immunofluorescent neurons throughout the brain of the brown anole (*Anolis sagrei*). This is therefore the first such study in a lizard. Anole lizards have been the subject of a multitude of studies examining the neural regulation of social behavior [e.g., 14,15,24,16–23], and CRF signaling may play a role in this behavioral regulation. Thus, increasing our understanding of the CRF distribution in lizards may eventually also provide insights into the regulation of lizard social behavior, a main focus of our research program.

In general, we find that brown anoles possess CRF distributions that are highly similar to that of other amniotes, especially those of other reptiles, and with a slightly more restricted distribution than that seen in mammals and birds. Although snakes lizards are closely related to snakes, and to a lesser degree to turtles [8], we do describe several notable differences between CRF distributions across these groups.

## Methods

### Subjects

This study examined the brains of three male brown anoles that were part of a previous study examining Ile^8^-oxytocin (mesotocin) colocalization with CRF in the hypothalamus [15]. Although we did not examine female brown anole brains, results from CRF studies in other reptiles have found no sex differences in intensity or distribution of signal [9–11]. Animals were housed in a terrarium (30.5 cm H x 26 cm W x 51 cm L) under long-day (14 h light:10 h dark) conditions. The terrarium was illuminated by a 40 W full-spectrum fluorescent light (Colortone 50, Philips, Amsterdam, Netherlands) suspended 20 cm above a wire-mesh terrarium lid. Heat was provided beyond ambient room temperature by means of a 60-W incandescent white light bulb suspended 5 cm above the terrarium lid. Reproductive status was verified at the time of sacrifice; all testes were determined to be enlarged from winter recrudescence. All procedures were approved by the Rhodes College Institutional Animal Care and Use Committee and are in accordance with federal guidelines.

### Immunohistochemistry

After subjects were euthanized, their brains were fresh dissected and submersion fixed overnight in 4% paraformaldehyde in 0.1 M phosphate buffer at 4°C, followed by cryoprotection with 30% sucrose in 0.1 M phosphate-buffered saline (PBS). Brains were frozen and sectioned into two 50-μm series, one of which was used for the present study. These sections were processed separately for rostral and caudal portions of the brain, using various antibodies [17]. We here briefly describe those antibodies pertinent to the present study. All sections were rinsed twice for 30 min in PBS, then blocked for 30 min in a solution of 2.5% donkey serum (Sigma-Aldrich), 0.3% Triton-X-100 (Fisher Scientific), and 0.05% sodium azide (Fluka) in PBS. Sections were then incubated for 40 h in the blocking solution with the addition of 0.1 µg/ml of guinea-pig polyclonal anti-CRF antibody (Bachem). Following two 30-min PBS rinses, the sections were incubated for 3 h in the blocking solution with the addition of 16 µg/ml of donkey anti-guinea pig Alexa Fluor 647 (Jackson ImmunoResearch). Following two more 30-min PBS rinses, the tissues were then mounted onto gelatin-coated slides, cleared with Xylene Substitute (Sigma-Aldrich) for 5 min, and coverslipped using Prolong Gold mounting medium with DAPI (Life Technologies). As reported more extensively in a previous study, the specificity of the CRF antibody in these tissues was demonstrated by preadsorption with 10X and 100X CRF peptide (Sigma-Aldrich), both of which eliminated CRF immunoreactivity [15].

### Image Analysis

Fluorescent photomicrographs of each brain section were obtained using an LSM 700 confocal microscope and Zen 2010 imaging software (Carl Zeiss Inc., Göttingen, Germany), using a 20X objective to capture a grid of images at 5 µm intervals. A maximum intensity projection of this stack was then created to produce a single two-dimensional photomicrograph, and the individual images were stitched together to generate a larger image of the entire brain section. This photomicrograph was exported using AxioVision 4.8 (Carl Zeiss Inc.) and analyzed in Adobe Photoshop CS6 (Adobe Systems Inc., San Jose, CA). Brightness and contrast were adjusted for clarity, but no further image manipulations were made. Brain regions were determined by reference to multiple atlases and publications [14,15,17,19,20,25–34].

## Results

CRF-immunoreactive cells and fibers were found throughout the brown anole brain (Fig. 1 and 2). Below, we describe CRF presence within specific neural locations.

**Figure 1.**
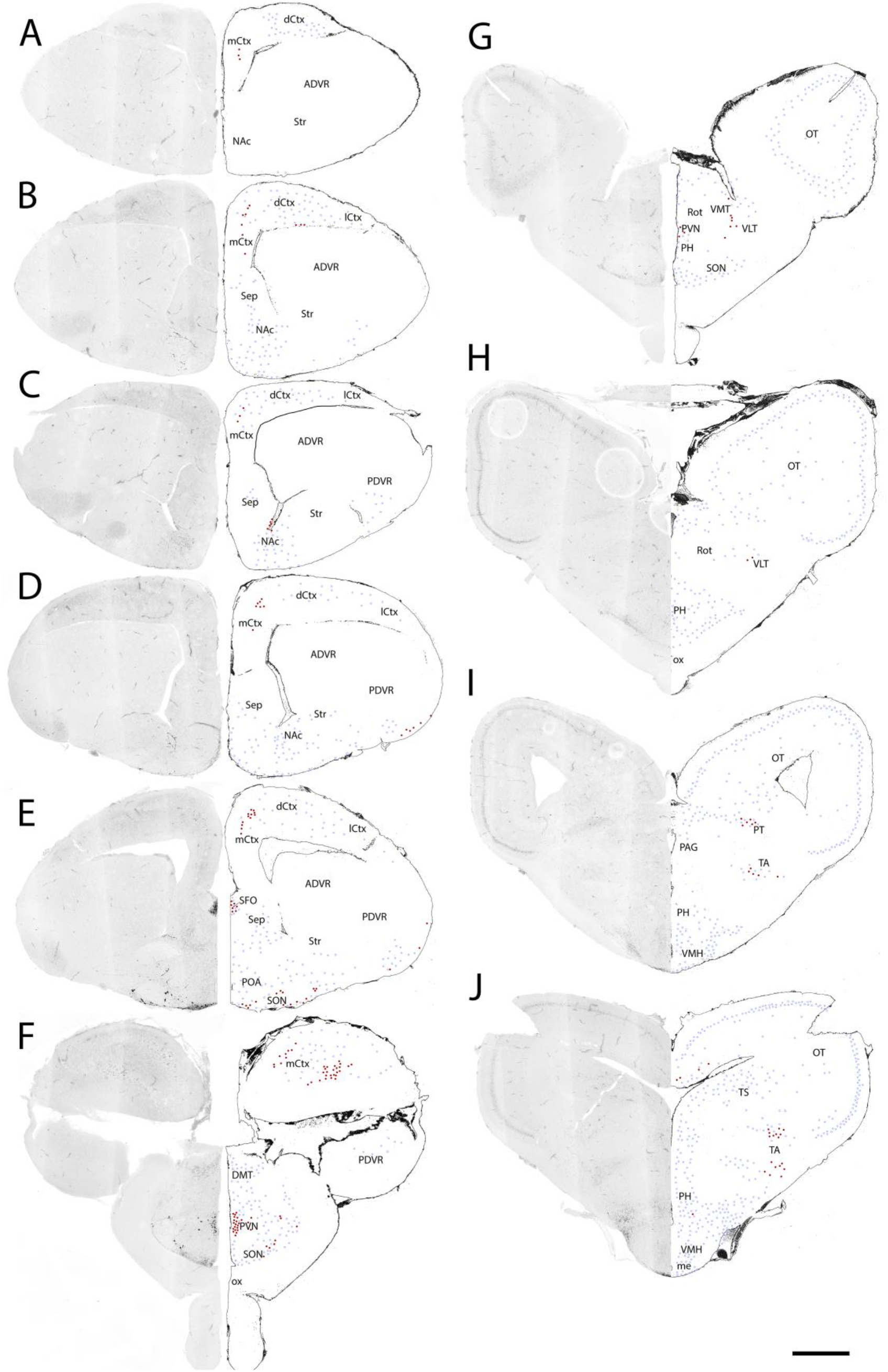
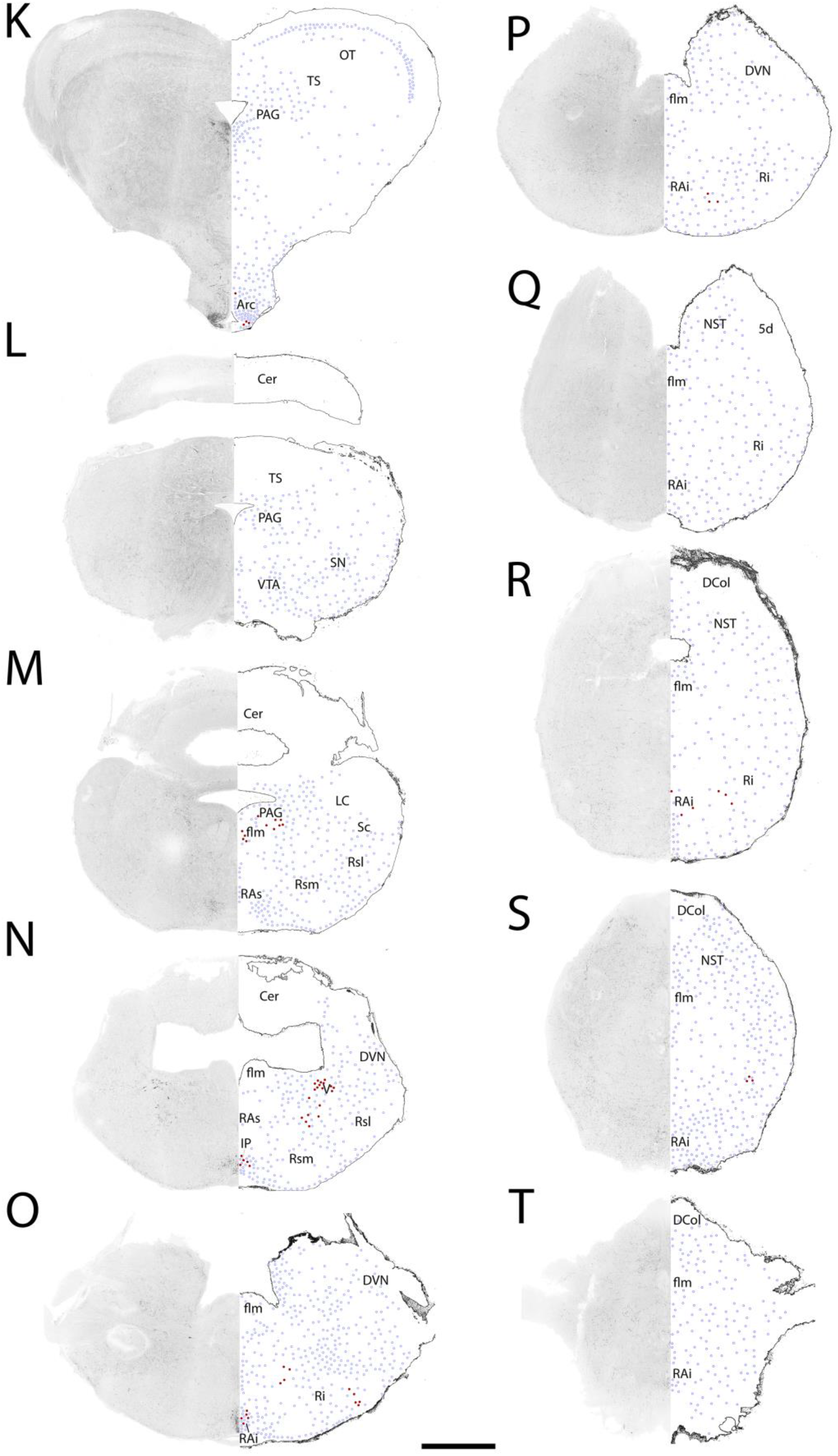
Representative sections from one brain, starting rostrally and progressing caudally. Sections are spaced 300 µm apart. Immunofluorescence in the left hemisphere is depicted, color-inverted, and a mirror image outline is present on the right (the single exception is panel N, which represents the right hemisphere due to damage to the left). Light blue circle outlines represent CRF-immunoreactive fibers, while dark red solid dots represent CRF-immunoreactive cells. The scale bar represents 500 µm. Abbreviations: 5d, descending tract of the trigeminal nerve; ADVR, anterior dorsal ventricular ridge; Arc, arcuate nucleus; Cer, cerebellum; DCol, nucleus of the dorsal column; dCtx, dorsal cortex; DMT, dorsomedial thalamic nucleus; DVN, dorsal vestibular nucleus; flm, medial longitudinal fasciculus; LC, locus coeruleus; lCtx, lateral cortex; mCtx, medial cortex; me, median eminence; Nac, nucleus accumbens; NST, nucleus of the solitary tract; OT, optic tectum; ox, optic chiasm; PAG, periaqueductal gray; PDVR, posterior dorsal ventricular ridge; PH, periventricular hypothalamic nucleus; POA, preoptic area; PON, paraventricular organ nucleus; PT, pretectal area; PVN, paraventricular nucleus of the hypothalamus; RAi, inferior raphe; RAs, superior raphe; Ri, inferior reticular nucleus; Rot, nucleus rotundus; Rsl, superior reticular nucleus, lateral; Rsm, superior reticular nucleus, medial; Sc, subcoeruleus; Sep, septum; SFO, subfornical organ; SN, substantia nigra; SON, supraoptic nucleus of the hypothalamus; Str, striatum; TA, tegmental area; TS, torus semicircularis; V, nucleus of the trigeminal nerve; VLT, ventrolateral thalamus; VMH, ventromedial hypothalamus; VMT, ventromedial thalamic nucleus; VTA, ventral tegmental area.

**Figure 2.**
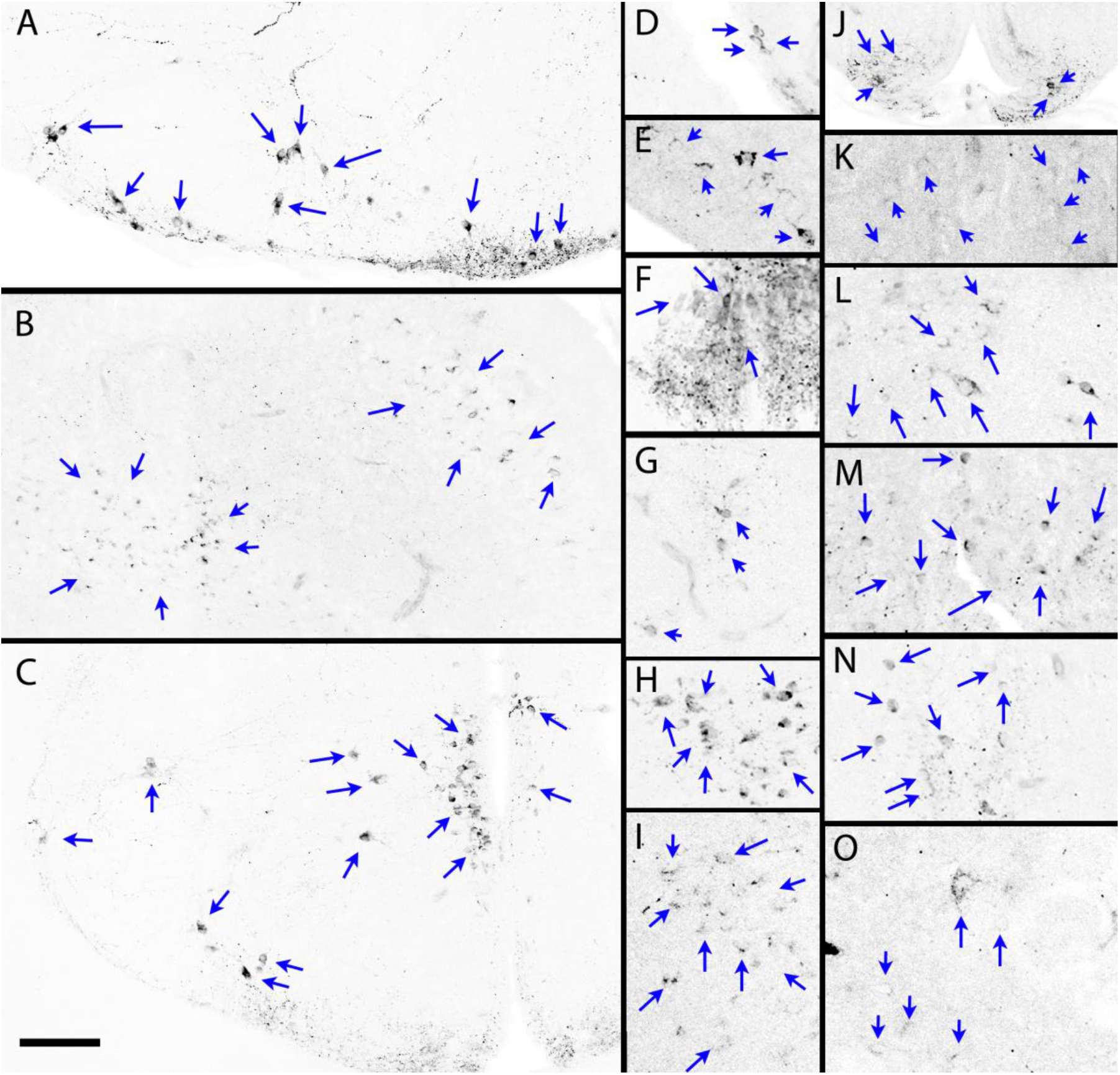
Color-inverted CRF immunofluorescent cells were present in select areas of the brown anole brain. Panels show: A, supraoptic nucleus (corresponds to Fig. 1E); B, medial cortex (corresponds to Fig. 1F); C, hypothalamic supraoptic and paraventricular nuclei (corresponds to Fig. 1F); D, nucleus accumbens (corresponds to Fig. 1C); E, ventral posterior dorsal ventricular ridge (ventral and lateral amygdalae; corresponds to Fig. 1D); F, subfornical nucleus (corresponds to Fig. 1E); G, lateral thalamus (corresponds to Fig. 1G); H, pretectal area (from one section caudal to Fig. 1I); I, midbrain tegmental area (corresponds to Fig. 1J); J, arcuate nucleus and medial eminence (corresponds to Fig. 1K); K, periaqueductal gray region, including oculomotor nerve nucleus (corresponds to Fig. 1M); L, nucleus of the trigeminal nerve (corresponds to Fig. 1N); M, interpeduncular nucleus (Fig. 1N); N, inferior raphe nucleus (corresponds to Fig. 1O); O, reticular nucleus (corresponds to Fig. 1O). Scale bar represents 100 µm for A-C and J, and 50 µm for the remaining panels.

### Forebrain

Most rostrally in the forebrain, CRF-immunoreactive fibers were present in the dorsal cortex. Moving caudally, the CRF fibers expanded throughout all other cortical regions, and also became apparent within the septum, striatum, the posterior dorsal ventricular ridge, the nucleus accumbens, and later also the preoptic area. CRF-immunoreactive cells were first apparent within the medial cortex, and more caudally within the dorsomedial, and dorsal cortices, as well as the nucleus accumbens. Later, prominent cell groups and heavy fiber innervation were found within the subfornical organ, supraoptic nucleus, and paraventricular nucleus of the hypothalamus. A small number of cells were also present in the ventral and ventrolateral posterior dorsal ventricular ridge, corresponding to the ventral and lateral amygdalae. CRF fibers were also present within various thalamic and hypothalamic nuclei. Moving further caudally, cells were present in the ventrolateral thalamus, as well as occasionally within the periventricular hypothalamus. Fibers were also present various hypothalamic regions, including the periventricular hypothalamus, and the ventromedial and ventrolateral hypothalamus. Finally, CRF cells and fibers were also present in the arcuate nucleus and the median eminence.

### Midbrain

Extensive CRF-immunoreactive fibers were present within the optic tectum, the periaqueductal gray area (also known as the central gray), the torus semicircularis, ventral tegmental area, and substantia nigra. Furthermore, fibers were present at the lateral aspects of the tegmentum. In the rostral midbrain, CRF-immunoreactive cells were located withing the pretectal area, the lateral tegmental area, and the torus semicircularis, dorsal to the fused third and optic ventricles. Cells were also present in caudal midbrain, in the area of the periaqueductal gray as well as the adjacent 3^rd^ nerve region. Finally, CRF cells were most caudally present within the interpeduncular nucleus and the nucleus of the trigeminal nerve.

### Hindbrain

Fibers were found extensively throughout the hindbrain, including in the raphe nuclei, reticular nuclei, dorsal vestibular nucleus, and nucleus of the solitary tract. Cells were present with the reticular nuclei, and inferior raphe.

## Discussion

Our results are generally consistent with the few other published central CRF distributions in reptiles, though with some notable exceptions. The other published reptile distributions comprise solely of a pair of snake species, the Brazilian pit viper *Bothrops jararaca* [9] and the viperine water snake *Natrix maura* [11], a study of the Caspian turtle *Mauremys capsica* [10], and two partial and redundant descriptions of the red-eared slider *Trachemys scripta* [12,13]. We predicted that the CRF distribution of brown anoles would more closely resemble that of snakes than turtles, as lizards and snakes share a much more recent common ancestor than either do with turtles [8].

Similar to what is seen in these other reptile species, brown anoles were found to possess CRF cells and fibers in the subfornical organ, paraventricular nucleus of the hypothalamus, and cortical regions [9–11]. Furthermore, like in three of the four other examined reptiles, brown anoles possessed cells and fibers in the nucleus accumbens (on the border of the ventral lateral septum), amygdaloid complex (ventral and lateral aspects of the posterior dorsal ventricular ridge), arcuate nucleus/infundibular recess/median eminence region, paraventricular organ nucleus, and reticular formation [9–13]. The location of cells within the amygdaloid complex varies across species, however, cells within the turtle *M. capsica* were in a location that is more similar to our findings, within posterior dorsal ventricular ridge[10], whereas those in the snake *N. maura* are shown as more likely within the medial amygdala region [11].

On the other hand, some of our findings were in direct contrast to what has been found in the other examined reptile species. Most strikingly, we found prominent CRF cells within the supraoptic nucleus whereas cells were found in neither of these regions in the snake and turtle studies, with solely fibers reported as present. Additionally, we found cell bodies in the lateral midbrain tegmentum, whereas only pretectal neurons were described in this region of the midbrain in the turtle species (and neither cells nor fibers in the snake species) [9–13].

In regard to other brain regions, there was a lack of consensus across the previously published reptile studies [9–13]. In some cases, our results lined up with those from the examined snake species, as expected, rather than the examined turtle species. For instance, as in brown anoles, both examined snake species possessed CRF cells within the trigeminal nerve and oculomotor nerve nuclei, while the turtle *M. capsica* did not (although it did possess cells slightly ventral to the oculomotor nerve nucleus, within the periaqueductal gray area). Similarly, our findings of CRF fibers within the septum, ventromedial hypothalamus, and habenula are in line with both snake studies, but not the *M. capsica* turtle study, which describes an absence of signal in these area (the *P. scripta* publications did not address these areas). However, at other times, our results surprisingly aligned more closely with those of the turtle studies than the snake studies. For instance, examination of *M. capsica* found CRF neurons within the interpeduncular nucleus/raphe region, while the examined snake species possess only fibers in this region; in the brown anole, we found cells in both the interpeduncular nucleus and inferior raphe, as well as fibers throughout this area. Furthermore, as mentioned above, both turtle species were found to possess CRF cells and fibers in the pretectal area, as in our study of brown anoles, while neither snake species was found to possess either cells or fibers in that region [9–13].

In other instances, species differences were present within taxonomic groupings, so our brown anole results were not strictly consistent with either snake or turtle findings, per se. For instance, CRF cells were described as present in the lamina terminalis in one of the examined snake species, *N. maura*, and one examined turtle species, *M. capsica*; however, in our study, we found only fibers in this region, much like in the other snake species, *B. jararaca* [9–11]. Additionally, we found CRF neurons in the lateral thalamus, and the turtle species *M. Capsica* was the only other reptile species with cells noted to be present in this area (described by the authors as the dorsolateral aggregation) [10].

When comparing our results to those in other amniotes, we generally see similarities of both mammals and birds with the brown anole brain. For instance, mice also possess CRF cells within cortical regions, amygdaloid regions, hypothalamic regions, and multiple hindbrain regions [1]. However, whereas they also produce CRF in neurons of the hypothalamic paraventricular nucleus, mice apparently do not possess CRF neurons within the supraoptic nucleus, instead producing the related urocortins 1 and 2 in this region [1]. However, CRF-producing neurons are present in the supraoptic nucleus of the rat [3,6] and tree shrew, *Tupaia belangeri* [4]. Other regions that match our CRF mapping include the arcuate nucleus and raphe nuclei, which possess harder to visualize CRF neurons that are more readily visible following colchicine-treatment in tree shrews [4]. Birds have also been found to possess widespread central CRF neurons [7]. Much like the cortical CRF populations in our study and other examined amniotes, birds also have widespread CRF neurons throughout their pallium [7]. Findings from chickens and Japanese quail mirror those in brown anoles by demonstrating the presence of CRF neurons in the supraoptic nucleus and along the third ventricle, especially within the hypothalamic paraventricular nucleus [7]. Furthermore, as in brown anoles, CRF cells are also present in periventricular hypothalamus, the amygdaloid nuclei, the lateral thalamus, the arcuate nucleus region, the pretectal nucleus, the optic tectum, the torus semicircularis, the periaqueductal gray area, and the raphe nuclei [7]. However, bird descriptions also noted the presence of CRF neurons with the locus coeruleus and subcoeruleus [11], while we found no neurons within this region. Birds also seem to possess generally more extensive hindbrain populations of CRF neurons than do brown anoles [11]. Both mammals [1,6] and birds [7] were also found to possess CRF neurons within the bed nucleus of the stria terminalis, a region where neurons have not been reported in our study, or in the other published reptile studies [9–13].

## Conclusions

The distribution of CRF-immunoreactive neurons within the brown anole brain is highly similar to that observed in snakes and turtles, the only other reptile species whose CRF distributions have been described. The main exception is the high presence of CRF neurons within the supraoptic nucleus of brown anoles, a region which has been described as containing only CRF fibers and not cells in other reptile species. The brown anole CRF distribution is furthermore fairly similar to, although more restricted than, that observed in mammals and birds. This is especially true for the absence of CRF neurons in the bed nucleus of the stria terminalis in the brains of brown anoles, and other reptiles, while this is a location of dense CRF neuron clustering in both mammals and birds.

## Acknowledgements

We would like to thank Rhodes College, as well as Dr. Charles and Mrs. Patricia Robertson, for their generous support.

